# Transcriptomics based prediction of metastasis in TNBC patients: Challenges in cross-platforms validation

**DOI:** 10.1101/2021.09.17.460812

**Authors:** Naorem Leimarembi Devi, Anjali Dhall, Sumeet Patiyal, Gajendra P. S. Raghava

## Abstract

Triple-negative breast cancer (TNBC) is more prone to metastasis and recurrence than other breast cancer subtypes. This study aimed to identify genes that can act as diagnostic biomarkers for predicting lymph node metastasis in TNBC patients. The transcriptomic data of TNBC with or without lymph node metastasis was acquired from TCGA, and the differentially expressed genes were identified. Further, logistic-regression method has been used to identify the top 15 genes (or 15 gene signatures) based on their ability to predict metastasis (AUC>0.65). These 15 gene signatures were used to develop machine learning techniques based prediction models; Gaussian Naïve Bayes classifier outperformed other with AUC>0.80 on both training and validation datasets. The best model failed drastically on nine independent microarray datasets obtained from GEO. We investigated the reason for the failure of our best model, and it was observed that the certain genes in 15 gene signatures were showing opposite regulating trends, i.e., genes are upregulated in TCGA-TNBC patients while it is downregulated on other microarray datasets or vice-versa. In conclusion, the 15 gene signatures may act as diagnostic markers for the detection of lymph node metastatic status in TCGA dataset, but quite challenging across multiple platforms. We also identified the prognostic potential of the 15 selected genes and found that overexpression of ZNRF2, FRZB, and TCEAL4 was associated with poor survival with HR>2.3 and p-value≤0.05. In order to provide services to the scientific community, we developed a webserver named “M_TNBC_Pred” for the prediction of metastatic and non-metastatic lymph node status of TNBC patients (http://webs.iiitd.edu.in/raghava/mtnbcpred/).

## Introduction

Over the past two decades, cancer is the leading cause of mortality in males and females globally. According to GLOBOCAN, in 2020, the worldwide cancer burden is estimated to be 19.3 million new cases and 10.0 million fatalities (Sung et al., 2021). Breast cancer has surpassed lung cancer to become the most frequently diagnosed cancer (Sung et al., 2021). An estimated 281,550 new breast cancer cases and 43,600 new deaths were reported in 2021 (Siegel et al., 2021a). Breast cancer can be divided into five subtypes based on gene expression profiling such as luminal A, luminal B, Human Epidermal Growth Factor Receptor 2 (HER2) over-expression, normal breast-like, and triple-negative breast cancer (TNBC) (Perou et al., 2000; Sorlie et al., 2001). TNBC accounts for 15–20% of all breast cancers, exhibits a high rate of relapse, increased metastatic potential, and poor prognosis. Due to the lack of target receptors such as Estrogen Receptor (ER), Progesterone Receptor (PR), and HER2 genes, TNBC patients do not get benefits from targeted therapies (Yao et al., 2017). The treatment options mostly rely on chemotherapy for early-stage and distant organ metastasis (Neophytou et al., 2018). TNBC patients tend to show lymph node metastasis at initial diagnosis and later spread to other organs like bone, lungs, brain, liver, etc (Garrido-Castro et al., 2019; Kennecke et al., 2010; Zhou et al., 2021). Generally, the lymphatic system helps maintain tissue fluid homeostasis, transporting antigen and immune cells to lymph nodes. However, disruption of this system will lead to severe diseases like lymphedema and cancer (Padera et al., 2016).

Previous studies have indicated that lymph node status is an important factor in determining breast cancer patients’ staging, prognosis, and overall survival (Al-Mahmood et al., 2018; Lee et al., 2021). At present, the surgical method to identify nodal status in TNBC includes axillary node dissection and sentinel node biopsy but causes various chronic side effects such as numbness and lymphedema (Abass et al., 2018). Therefore, it is required to search for more accurate markers for the preoperative identification of lymph node metastasis (LNM) to reduce the mortality rate for TNBC patients. Recent advances in high-throughput technologies generate a massive amount of transcriptomic and genomic public repositories such as The Cancer Genome Atlas (TCGA) and Gene Expression Omnibus (GEO) (Cancer Genome Atlas Research Network et al., 2013; Edgar, 2002). The genomic data provide gainful insights to understand the biological nature of different malignancies. Recently, several studies utilized genomic data to identify potential biomarkers in metastatic cancer patients (Mathe et al., 2015; Meyerson et al., 2010; Zhu et al., 2019). Thus, the prediction or diagnosis of lymph node metastasis is essential for TNBC patients.

In this study, we have attempted to develop models based on the mRNA expression/ transcriptomic profiles to classify lymph node metastatic and non-metastatic patients in TNBC. We identified 15 gene panels that can act as diagnostic biomarkers for the detection of LNM-TNBC. Additionally, survival analysis was performed using the expression and clinical data of TCGA-TNBC patients, to stratify high-risk and low-risk groups. We have also attempted to validate the gene signatures across a variety of platforms obtained from GEO. Finally, we have developed a webserver, “M_TNBC_Pred”, based on the gene expression data of TNBC patients that could successfully distinguish metastatic/ non-metastatic TNBC samples.

## Materials and Methods

### Dataset Collection

The level 3 mRNA read count data for TCGA breast cancer samples were retrieved from Broad GDAC Firehose (http://gdac.broadinstitute.org/). The clinical information such as ER-, PR-, HER2- defined according to immunohistochemically staining and fluorescence in situ hybridization results were filtered (Dent et al., 2007; Hammond et al., 2010; Wolff et al., 2007). 157 samples were obtained consisting of 53 TNBC LNM and 104 TNBC without LNM. As well as different microarray datasets were obtained from GEO such as GSE20685 (Kao et al., 2011), GSE19697 (Lin et al., 2010), GSE32646 (Miyake et al., 2012), GSE20711 (Dedeurwaerder et al., 2011), GSE58812 (Jézéquel et al., 2015), GSE76275 (Burstein et al., 2015; den Hollander et al., 2016), GSE31448 (Sabatier et al., 2011) of Affymetrix platform (GPL570), GSE46581 (Lindner et al., 2013) of Illumina (GPL8432) and GSE40206 (Bashir et al., 2015) of Agilent (GPL4133) for independent validation. The distribution of metastatic and non-metastatic LN-TNBC samples across different datasets are shown in Figure 1.

**Figure 1:**
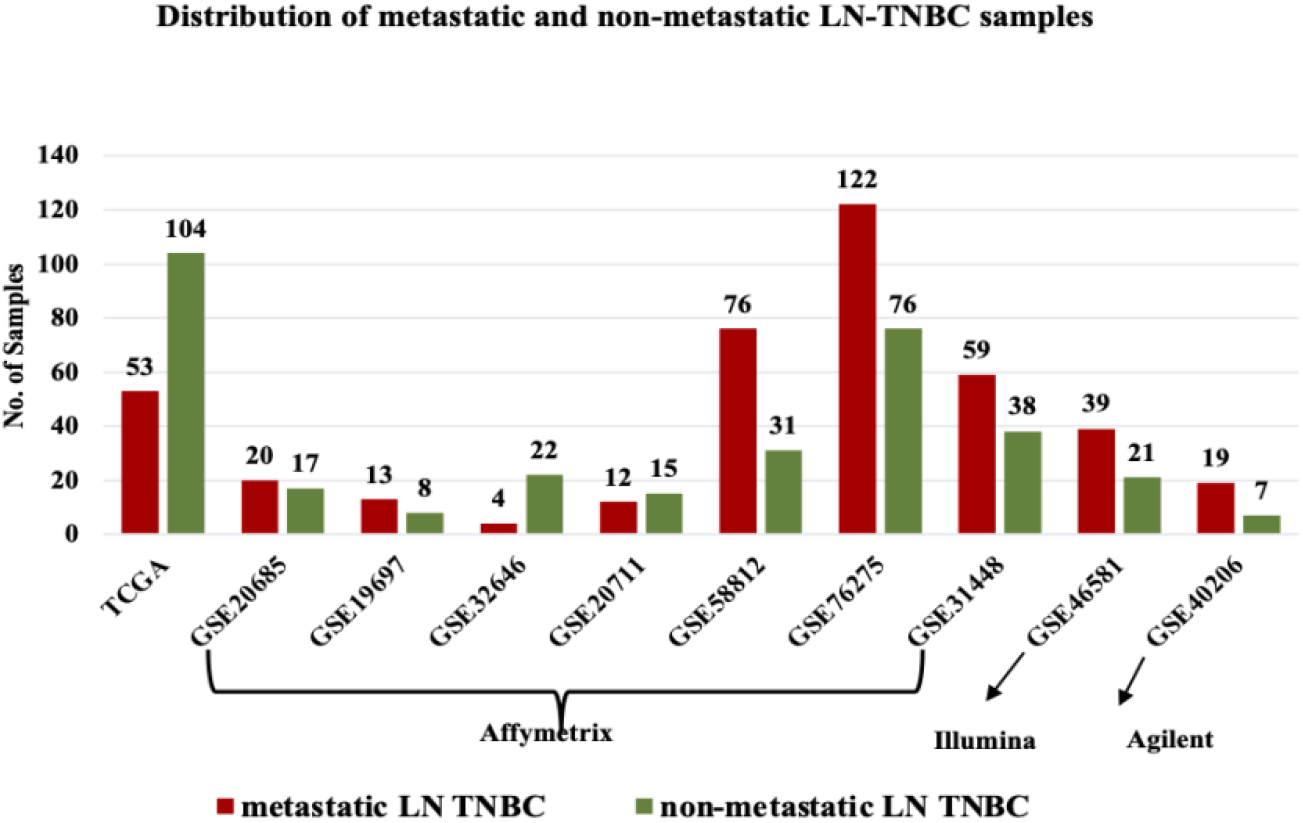
Distribution of metastatic and non-metastatic LN-TNBC patients in different platforms

### Dataset Pre-processing

The read counts were normalized to counts per million (CPM) using edgeR TMM-method (Robinson et al., 2010). Then, CPM was transformed to log2-CPM using voom() in limma (Ritchie et al., 2015). In case of microarray datasets, with Affymetrix platform, CEL files were pre-processed with background correction, quantile normalization and log2 transformation using affy package in R (Gautier et al., 2004). For Illumina and Agilent dataset, raw files were normalized using limma package, which performs normexp background correction, quantile normalised and log2 transformation (Ritchie et al., 2015). Finally, the average of several probes matching a single gene was calculated using python script for each dataset. These microarray datasets are used as an external validation dataset.

### Differential Expression Analysis

The normalized gene expression data was utilised for differentially expression analysis. Student’s *t*-test such as Welch *t*-test and Wilcoxon *t*-test was used to identify the differentially expressed genes (DEGs) in metastatic and non-metastatic LN-TNBC patients. It is implemented using in-house R scripts after samples were assigned to the respective class (Kaur et al., 2019). Thus, a set of genes was selected that showed statistically differentially expression between two classes of samples using Bonferroni adjusted p-value<0.05. The genes having significant positive mean-difference were considered as upregulated genes. On the other hand, the genes negative mean difference as downregulated genes.

### Feature Ranking

Feature selection is a critical phase in the machine learning process (Guyon, Isabelle, 2003). Thus, the ranking was performed using the logistic-regression method (Tolles & Meurer, 2016). In this method, the model was developed on each feature using logistic regression classifier. The features or genes were ranked based on their performance (AUC) to predict metastasis in TNBC.

### Classification Algorithms

In this study, prediction models were developed to distinguish metastatic and non-metastatic LN-TNBC based on the identified features using several machine learning algorithms. It includes Logistic Regression (LR), Gaussian Naive Bayes (GNB), eXtreme Gradient Boosting (XGB), Decision Tree (DT), Random Forest (RF), Support Vector Classifier (SVC), and K-Nearest Neighbors (KNN). The algorithms were implemented using the sklearn package in Python (Pedregosa, F. et al, 2011).

### Cross-validation Technique

To assess the performance of generated prediction models, the 5-fold cross-validation technique was implemented (Dhall et al., 2020). Previous studies employed an 80:20 ratio to split the dataset into training (80%) and validation dataset (20%) (Bhalla et al., 2019; Dhall et al., 2021; Sharma et al., 2021). Thus, we used this standard protocol and divided the main dataset into 80:20 ratio. Here, the 80% training data was split into 5-fold, of which four sets were considered as training and the remaining fifth set as testing. This process was repeated five times so that each set was used for training and testing. Then, the final performance of the model is the average of all five sets evaluated on 20% validation dataset (Dhall et al., 2021; Patiyal et al., 2020; Sharma et al., 2021).

### Performance Metrics

The performance measures of each classification model developed in this study were computed using threshold-independent and threshold-dependent parameters. In the case of threshold dependent parameters, four statistical measures such as sensitivity, specificity, accuracy, and MCC were computed using the following equations:

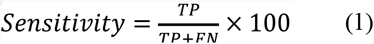

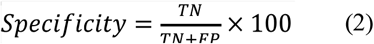

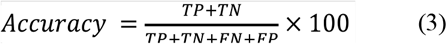

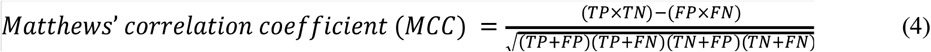

Where *TP*: true positive, *FN*: false negative, *FP*: false positive and *TN*: true negative, respectively. Also, the threshold-independent parameter, i.e., area under the receiver operating characteristic curve (AUC), was calculated to assess the model performance.

### Visualization of Samples

t-Distributed Stochastic Neighbor Embedding (t-SNE) is a nonlinear dimensionality reduction technique used to visualize the high-dimensional data. Using this technique, the distribution of samples was visualized based on selected 15 features to check if data-sets are segregating into defined classes. The visualization was performed using R packages such as Rtsne and scatter-plot3d (Van Der Maaten, 2014).

### Survival Analysis

To measure the prognostic potential of the top-15 genes, we have performed univariate survival analysis using the survival package in R (V.3.5.1) (Therneau, 2014). The clinical information like tumor stage, lymph node status, age, overall survival (OS) time, and vital status of 157 and 107 LN-TNBC samples were collected from TCGA and GSE58812, respectively (Supplementary Table S1). The dataset was split into two groups (i.e., high-risk and low-risk) based on the mean expression value of the genes. To compute the survival significance of two groups, a log-rank test was used. In addition, we have reported hazard ratio (HR), 95% confidence interval (CI), p-value and concordance. For graphical visualization of high-risk and low-risk groups, we have provided the Kaplan-Meier (KM) survival curves (Goel et al., 2010).

## Results

The mRNA expression data of 20,531 genes for LNM-TNBC samples were obtained from TCGA. Initially, the VarianceThreshold method of scikit learn library was implemented for filtering out low variance genes (Pedregosa, Fabian, 2011). After removing the low variance genes, we were left with 17297 genes. Further, a total of 1010 DEGs, including 643 upregulated and 367 downregulated genes, were identified in metastatic and non-metastatic LN-TNBC samples (Supplementary Table S2).

### Single Gene Biomarker

The logistic-regression method was used to rank the DEGs. The performance of all 1010 DEGs was calculated as mentioned in Supplementary Table S3. The top-15 genes were selected having an AUC value greater than 0.65 (Table 1). The expression patterns of top-15 genes were represented as a boxplot. Here, the red color indicates metastatic and green as non-metastatic LN-TNBC (Figure 2). Accordingly, DHRS7, BAIAP3, ZNRF2, ETFDH, HBG1, RIOK2, TCEAL4, TCF21, and FRZB genes were significantly upregulated in LN-TNBC patients. On the other hand, POU4F1, COL24A1, TRPA1, IBSP, VIL1, and PSAT1 genes were significantly downregulated in LN-TNBC samples.

**Table 1:** Performance of top 15 genes on TCGA LN-TNBC dataset.

| Feature | Sensitivity | Specificity | AUC | Accuracy | MCC |
| --- | --- | --- | --- | --- | --- |
| DHRS7 | 67.925 | 68.269 | 0.689 | 68.153 | 0.345 |
| ZNRF2 | 62.264 | 66.346 | 0.689 | 64.968 | 0.273 |
| VIL1 | 64.151 | 63.462 | 0.683 | 63.694 | 0.262 |
| BAIAP3 | 64.151 | 61.538 | 0.677 | 62.42 | 0.243 |
| PSAT1 | 66.038 | 67.308 | 0.674 | 66.879 | 0.318 |
| RIOK2 | 64.151 | 62.5 | 0.671 | 63.057 | 0.253 |
| ETFDH | 62.264 | 63.462 | 0.669 | 63.057 | 0.244 |
| HBG1 | 60.377 | 64.423 | 0.666 | 63.057 | 0.236 |
| TCF21 | 64.151 | 62.5 | 0.664 | 63.057 | 0.253 |
| FRZB | 66.038 | 68.269 | 0.663 | 67.516 | 0.327 |
| TCEAL4 | 62.264 | 60.577 | 0.661 | 61.146 | 0.216 |
| IBSP | 64.151 | 62.5 | 0.66 | 63.057 | 0.253 |
| TRPA1 | 62.264 | 61.538 | 0.658 | 61.783 | 0.226 |
| COL24A1 | 62.264 | 64.423 | 0.657 | 63.694 | 0.254 |
| POU4F1 | 60.377 | 60.577 | 0.656 | 60.51 | 0.199 |

**Figure 2:**
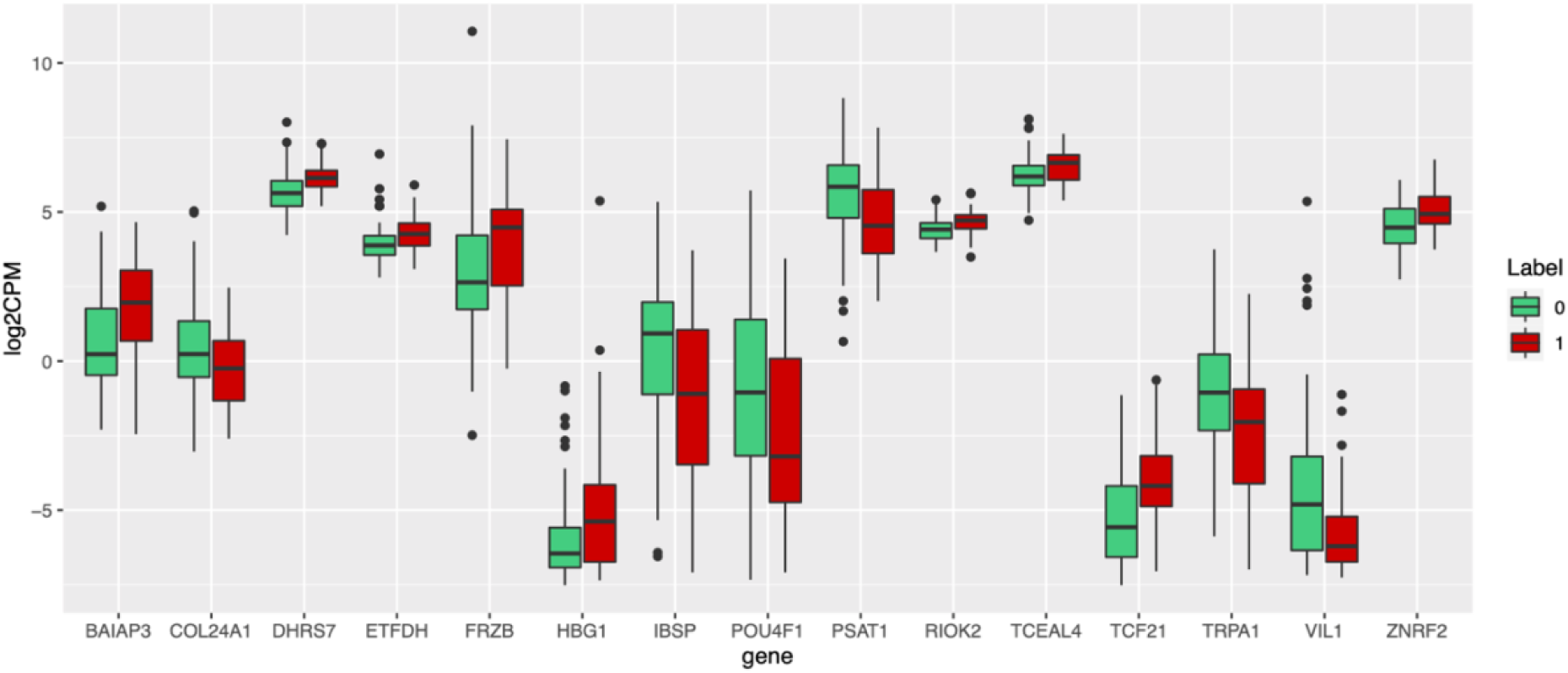
Box plot representing the expression pattern of top-15 genes in the TCGA-TNBC dataset, where red color indicates the metastatic and green with non-metastasis LN-TNBC

Genes, namely DHRS7 and ZNRF2, achieved the highest performance in classification with an AUC of 0.689 with balanced sensitivity and specificity. Therefore, DHRS7 and ZNRF2 may act as potential biomarkers for LN-TNBC.

### Multiple Gene-based Biomarkers

Next, we have used the top-15 genes to develop different classification models implemented by GNB, RF, LR, DT, KNN, XGB, and SVC. We compute the performance of top-2, 3, 4…,15 genes as mentioned in Supplementary Table S4. We observed that the models based on top-5 genes (DHRS7, ZNRF2, VIL1, BAIAP3, PSAT1) perform quite well with an AUC of 0.76 on training and validation dataset as provided in Supplementary Table S4. The GNB-based model based on top-15 genes outperformed the other methods and achieved a maximum AUC of 0.82 and 0.81 on training and validation datasets, respectively, with balanced specificity and sensitivity (Table 2). From the above results, we have observed that 15 mRNA expression-based features are able to classify metastatic and non-metastatic LN-TNBC samples.

**Table 2:** Performance of machine-learning models on training and validation dataset with top 15 genes.

| Classifier | Training Dataset |  |  |  |  | Validation Dataset |  |  |  |  |
| --- | --- | --- | --- | --- | --- | --- | --- | --- | --- | --- |
|  | Sensitivity | Specificity | Accuracy | AUC | MCC | Sensitivity | Specificity | Accuracy | AUC | MCC |
| **GNB** | 75.56 | 76.25 | 76.00 | 0.82 | 0.50 | 75.00 | 79.17 | 78.12 | 0.81 | 0.50 |
| **LR** | 77.78 | 75.00 | 76.00 | 0.80 | 0.51 | 75.00 | 70.83 | 71.88 | 0.81 | 0.40 |
| **XGB** | 73.33 | 71.25 | 72.00 | 0.80 | 0.43 | 50.00 | 66.67 | 62.50 | 0.64 | 0.15 |
| **DT** | 57.78 | 56.25 | 56.80 | 0.59 | 0.14 | 50.00 | 54.17 | 53.12 | 0.47 | 0.04 |
| **SVC** | 77.78 | 76.25 | 76.80 | 0.81 | 0.52 | 75.00 | 62.50 | 65.62 | 0.78 | 0.32 |
| **RF** | 77.78 | 73.75 | 75.20 | 0.82 | 0.50 | 75.00 | 75.00 | 75.00 | 0.75 | 0.45 |
| **KNN** | 77.78 | 80.00 | 79.20 | 0.82 | 0.56 | 75.00 | 70.83 | 71.88 | 0.77 | 0.40 |

GNB: Gaussian Naïve Bayes, LR: Logistic Regression, XGB: eXtreme Gradient Boosting, DT: Decision Tree, SVC: Support Vector Classifier, RF: Random Forest, KNN: K-Nearest Neighbours, AUC: area under the receiver operating characteristic curve, MCC: Matthews' correlation coefficient

Further, we visualized the samples based on the expression profiles of these genes by utilizing the t-SNE. As shown in Figure 3, the metastatic samples stratify the non-metastatic group based on the top-15 genes, with a few overlapping samples.

**Figure 3:**
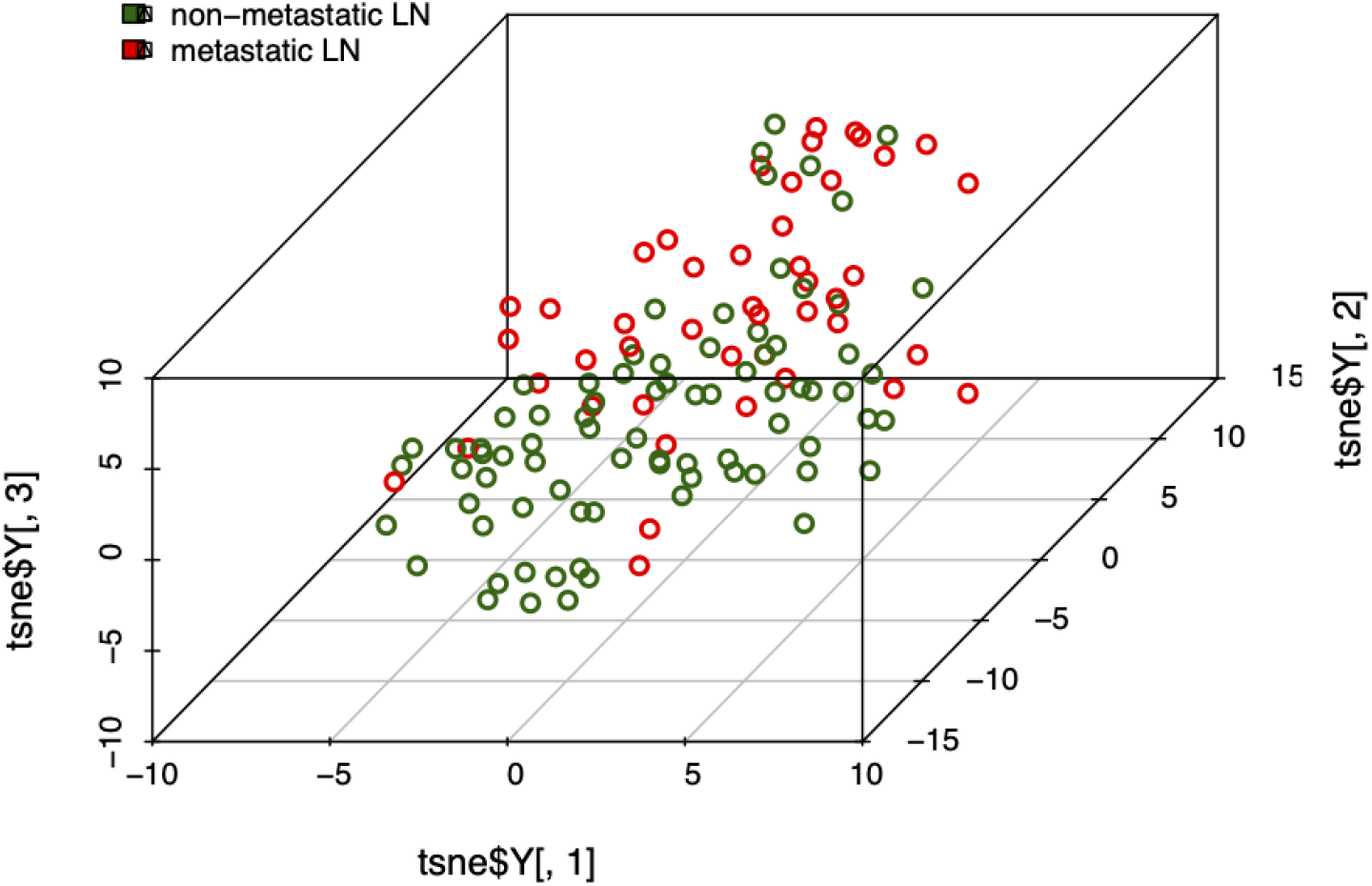
t-SNE visualization of metastatic and non-metastatic LN-TNBC based on top-15 selected features

### Interaction Network and Enrichment Analysis

Interaction network was generated using GeneMANIA (http://genemania.org/) (Franz et al., 2018) and visualised in Cytoscape (Shannon et al., 2003) to gain insights into the underlying molecular links of top-15 genes. The network was constructed based on multiple layers of existing evidence that includes the genetic interaction accounts for 7.36%, co-expression for 73.90%, and co-localization for 18.73%. The co-expression links account for the largest proportion, which demonstrated the presence of highly biological interactions among these genes (Figure 4A). Besides this, enrichment analysis was done using Enrichr (Kuleshov et al., 2016) on the selected genes. GO analysis revealed that the DEGs were enriched in processes such as positive regulation of neurotransmitter transport, lung morphogenesis, cell differentiation etc. KEGG pathway enrichment analysis demonstrated that the DEGs were mainly associated with serine metabolism, Vitamin B6 metabolism, etc. Further, Wiki Pathway indicates that the genes are involved in Interleukin-11 signaling pathway, osteoblast signaling, etc (Figure 4B).

**Figure 4:**
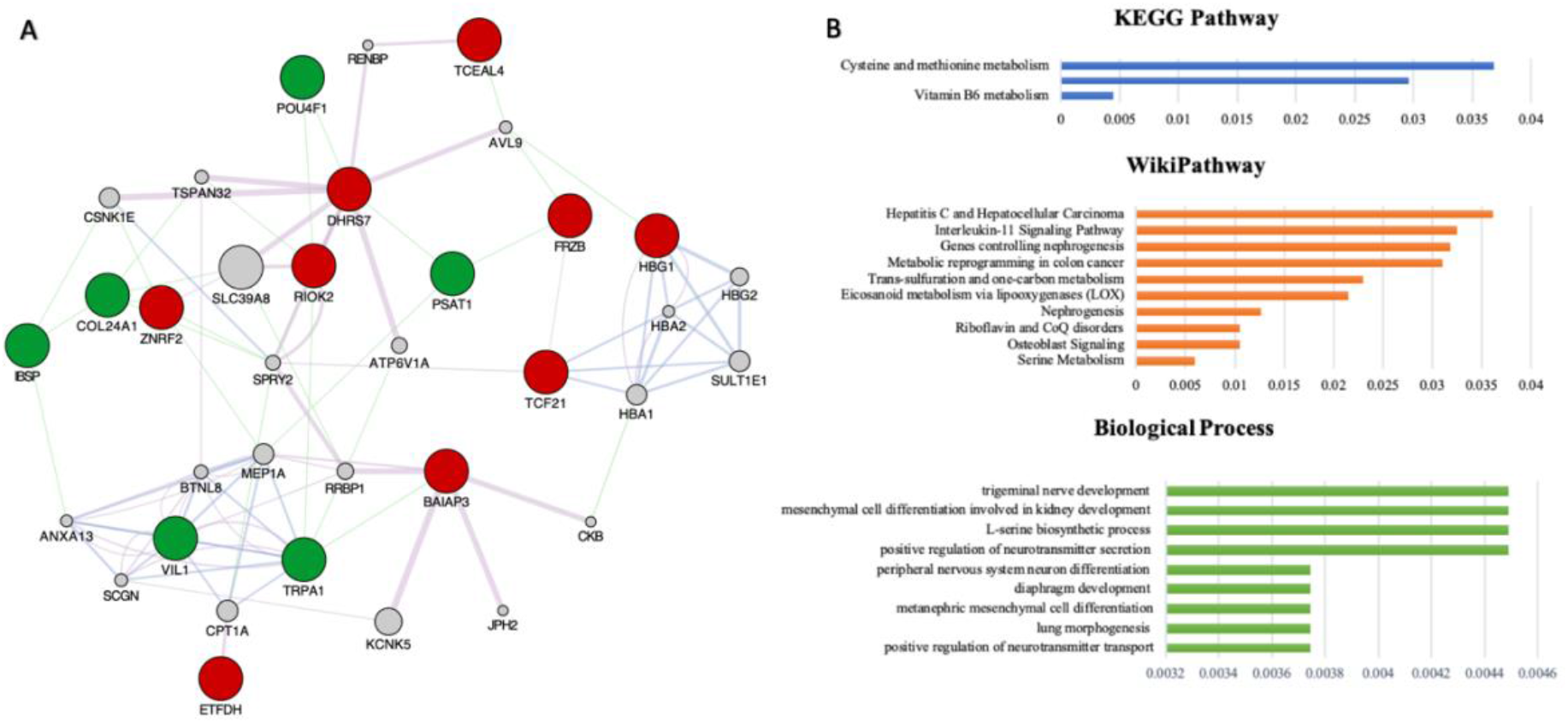
A) Interaction network where the red and green nodes are the input genes, and the grey nodes are the predicted nodes. The violet link is the co-expression interaction, the blue link indicates co-localization, and the green is the genetic interaction. B) Pathway and Biological process enrichment analysis of the identified genes

### Cross-platform Validation

The performance of different independent microarray datasets (GSE20685, GSE19697, GSE32646, GSE20711, GSE58812, GSE76275, GSE31448, GSE46581, and GSE40206) is evaluated on the parameters of the GNB-based model developed on 15 genes. As shown in Table 3, we observed that the 15-genes based model performed quite well on TCGA validation dataset; however, it performs poorly on another cross-validation dataset. The dataset GSE40206 of Agilent platform is able to achieve a maximum AUC of 0.60 with balanced sensitivity and specificity, as mentioned in (Table 3).

**Table 3:**
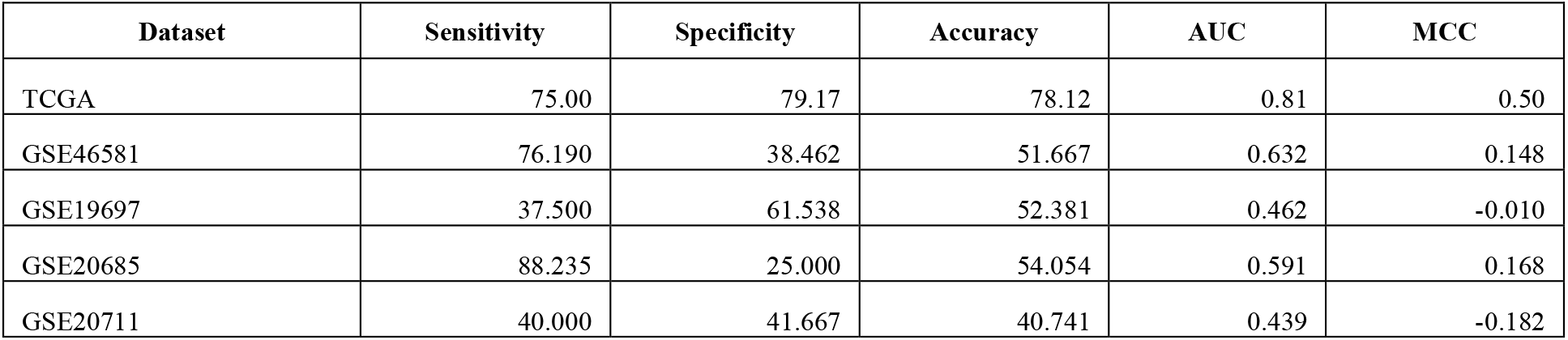
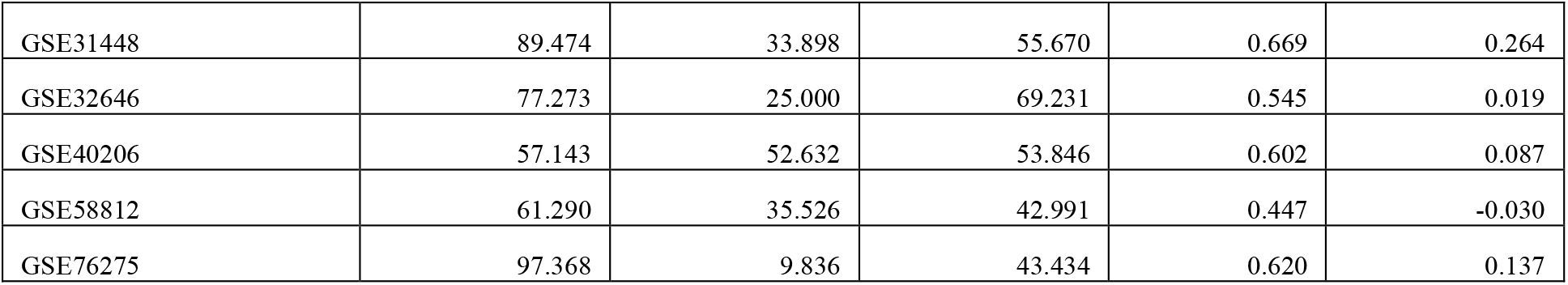
Performance of best GNB-model on independent microarray datasets with top-15 genes.

| Dataset | Sensitivity | Specificity | Accuracy | AUC | MCC |
| --- | --- | --- | --- | --- | --- |
| TCGA | 75.00 | 79.17 | 78.12 | 0.81 | 0.50 |
| GSE46581 | 76.190 | 38.462 | 51.667 | 0.632 | 0.148 |
| GSE19697 | 37.500 | 61.538 | 52.381 | 0.462 | -0.010 |
| GSE20685 | 88.235 | 25.000 | 54.054 | 0.591 | 0.168 |
| GSE20711 | 40.000 | 41.667 | 40.741 | 0.439 | -0.182 |
| GSE31448 | 89.474 | 33.898 | 55.670 | 0.669 | 0.264 |
| GSE32646 | 77.273 | 25.000 | 69.231 | 0.545 | 0.019 |
| GSE40206 | 57.143 | 52.632 | 53.846 | 0.602 | 0.087 |
| GSE58812 | 61.290 | 35.526 | 42.991 | 0.447 | -0.030 |
| GSE76275 | 97.368 | 9.836 | 43.434 | 0.620 | 0.137 |

### Robustness of Signature Genes

The robustness of the gene signatures was evaluated on nine independent microarray datasets (i.e., 7 Affymetrix, 1 Illumina, and 1 Agilent). The differential expression levels of 15 genes were identified in between the metastasis and non-metastasis TNBC patients. From this analysis, we observed that these genes show opposite regulating patterns, i.e., genes are upregulated in TCGA-TNBC patients while they are downregulated on other microarray datasets or vice-versa, as shown in Table 4.

**Table 4:**
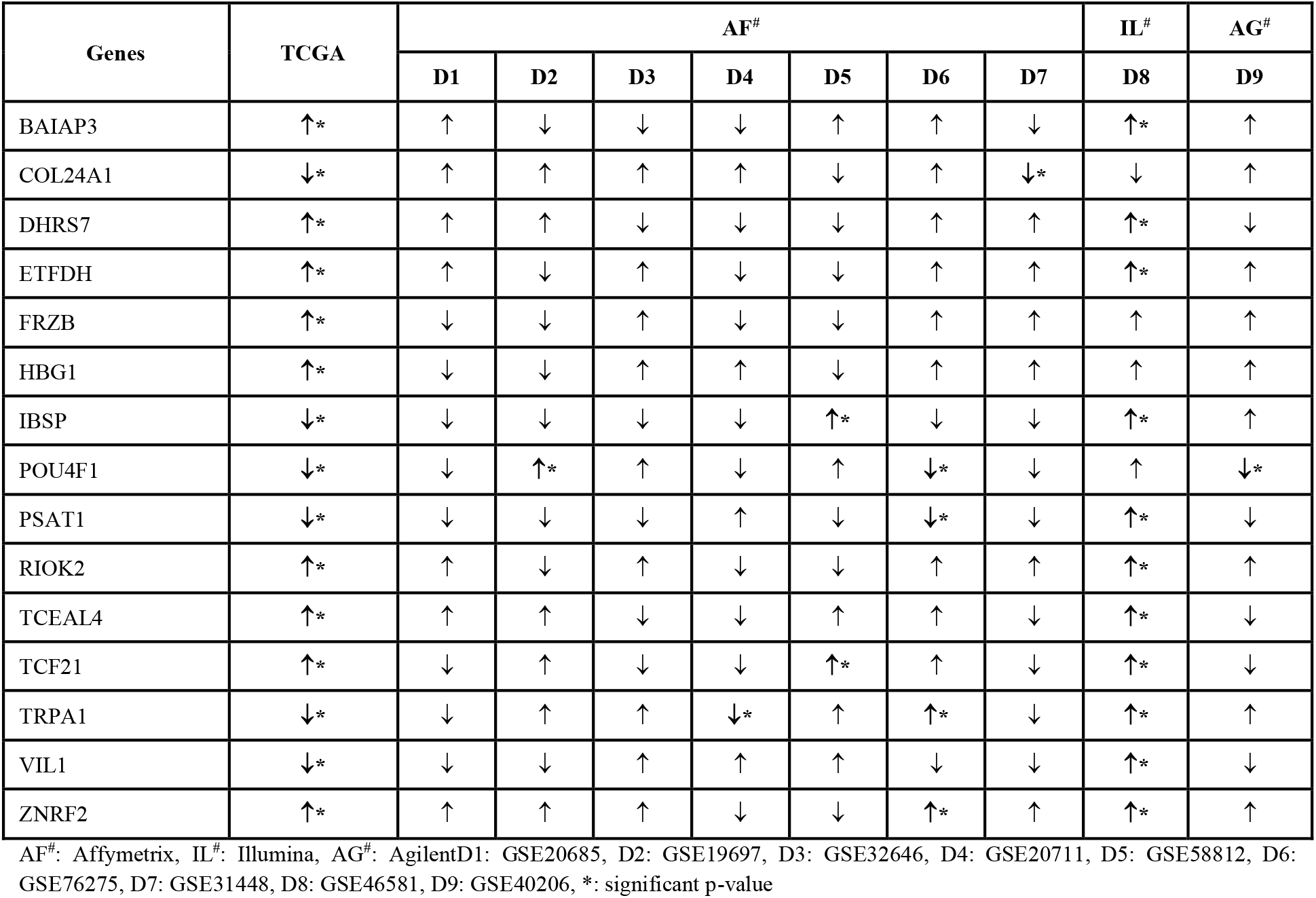
Differential expression analysis of the 15 genes with their regulating trends based on mean-difference on different datasets

| Genes | TCGA | AF<sup>#</sup> |  |  |  |  |  |  | IL<sup>#</sup> | AG<sup>#</sup> |
| --- | --- | --- | --- | --- | --- | --- | --- | --- | --- | --- |
|  |  | D1 | D2 | D3 | D4 | D5 | D6 | D7 | D8 | D9 |
| BAIAP3 | ↑* | ↑ | ↓ | ↓ | ↓ | ↑ | ↑ | ↓ | ↑* | ↑ |
| COL24A1 | ↓* | ↑ | ↑ | ↑ | ↑ | ↓ | ↑ | ↓* | ↓ | ↑ |
| DHRS7 | ↑* | ↑ | ↑ | ↓ | ↓ | ↓ | ↑ | ↑ | ↑* | ↓ |
| ETFDH | ↑* | ↑ | ↓ | ↑ | ↓ | ↓ | ↑ | ↑ | ↑* | ↑ |
| FRZB | ↑* | ↓ | ↓ | ↑ | ↓ | ↓ | ↑ | ↑ | ↑ | ↑ |
| HBG1 | ↑* | ↓ | ↓ | ↑ | ↑ | ↓ | ↑ | ↑ | ↑ | ↑ |
| IBSP | ↓* | ↓ | ↓ | ↓ | ↓ | ↑* | ↓ | ↓ | ↑* | ↑ |
| POU4F1 | ↓* | ↓ | ↑* | ↑ | ↓ | ↑ | ↓* | ↓ | ↑ | ↓* |
| PSAT1 | ↓* | ↓ | ↓ | ↓ | ↑ | ↓ | ↓* | ↓ | ↑* | ↓ |
| RIOK2 | ↑* | ↑ | ↓ | ↑ | ↓ | ↓ | ↑ | ↑ | ↑* | ↑ |
| TCEAL4 | ↑* | ↑ | ↑ | ↓ | ↓ | ↑ | ↑ | ↓ | ↑* | ↓ |
| TCF21 | ↑* | ↓ | ↑ | ↓ | ↓ | ↑* | ↑ | ↓ | ↑* | ↓ |
| TRPA1 | ↓* | ↓ | ↑ | ↑ | ↓* | ↑ | ↑* | ↓ | ↑* | ↑ |
| VIL1 | ↓* | ↓ | ↓ | ↑ | ↑ | ↑ | ↓ | ↓ | ↑* | ↓ |
| ZNRF2 | ↑* | ↑ | ↑ | ↑ | ↓ | ↓ | ↑* | ↑ | ↑* | ↑ |

AF<sup>#</sup>: Affymetrix, IL<sup>#</sup>: Illumina, AG<sup>#</sup>: AgilentD1: GSE20685, D2: GSE19697, D3: GSE32646, D4: GSE20711, D5: GSE58812, D6: GSE76275, D7: GSE31448, D8: GSE46581, D9: GSE40206, \*: significant p-value

### Risk Assessment of Signature Genes

From univariate survival analysis, we observed that 6 genes (3-upregulated and 3-downregulated) significantly stratify high-risk and low-risk groups. Interestingly, we found that high expression of ZNRF2 [HR = 2.711, p-value = 0.03], FRZB [HR = 2.395, p-value = 0.05] and TCEAL4 [HR=2.254, p-value =0.05] were associated with a poor survival for metastatic group. Whereas, the downregulation of PSAT1 [HR = 0.41, p-value = 0.04], TRPA1 [HR = 0.329, p-value = 0.01] and VIL1 [HR = 0.217, p-value = 0.013] genes is associated with better overall survival, as depicted in Figure 5. The complete results of survival analysis with HR, 95% Cl, p-value, concordance is given in Table 5. The robustness of the prognostic signature was evaluated in the external microarray dataset (GSE58812), having clinical information of TNBC patients. We found that higher expression of two genes ZNRF2 [HR=2.09, p-value=0.05] and FRZB [HR=2.74, p-value=0.02] is significantly associated with the poor prognosis of the patients (Table 6).

**Figure 5:**
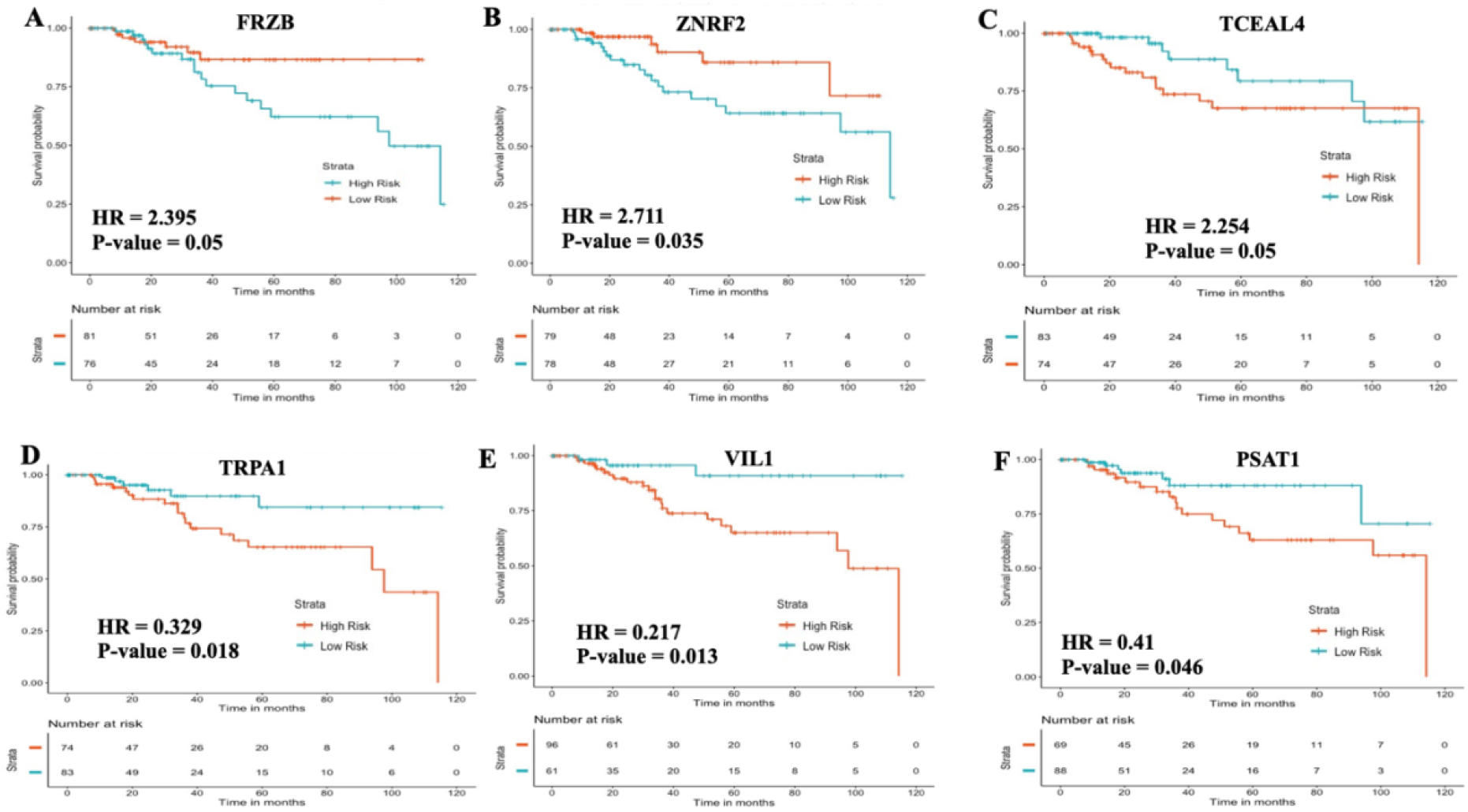
Kaplan Meier survival curves for risk estimation in LN-TNBC TCGA cohort based on the expression data where, (A) FRZB, (B) ZNRF2, (C) TCEAL4, are bad prognostic markers and associated with poor survival, whereas (D) TRPA1, (E) VIL1, and (F) PSAT1, are good prognostic markers and associated with better survival

**Table 5:** Survival analysis to stratify TNBC patients low-risk and high-risk groups on training dataset

| Gene | #G1 | #G2 | HR | p-Value | 95% CI | Concordance | Log (p-value) |
| --- | --- | --- | --- | --- | --- | --- | --- |
| BAIAP3 | 85 | 72 | 1.211 | 0.635 | 0.55-2.66 | 0.492 | 0.635 |
| COL24A1 | 87 | 70 | 1.318 | 0.502 | 0.58-2.95 | 0.497 | 0.504 |
| DHRS7 | 79 | 78 | 1.606 | 0.246 | 0.72-3.57 | 0.564 | 0.241 |
| ETFDH | 87 | 70 | 1.618 | 0.246 | 0.71-3.64 | 0.555 | 0.242 |
| FRZB | 81 | 76 | 2.394 | 0.053* | 0.98-5.79 | 0.562 | 0.04* |
| HBG1 | 103 | 54 | 0.822 | 0.646 | 0.35-1.89 | 0.559 | 0.643 |
| IBSP | 66 | 91 | 0.989 | 0.978 | 0.44-2.22 | 0.498 | 0.978 |
| POU4F1 | 80 | 77 | 0.451 | 0.075 | 0.18-1.08 | 0.619 | 0.062 |
| PSAT1 | 88 | 69 | 0.41 | 0.046* | 0.17-0.98 | 0.596 | 0.036* |
| RIOK2 | 82 | 75 | 1.944 | 0.125 | 0.83-4.54 | 0.609 | 0.115 |
| TCEAL4 | 83 | 74 | 2.254 | 0.050* | 0.97-5.23 | 0.655 | 0.050* |
| TCF21 | 81 | 76 | 2.132 | 0.078 | 0.91-4.94 | 0.581 | 0.068 |
| TRPA1 | 83 | 74 | 0.329 | 0.018* | 0.13-0.82 | 0.605 | 0.011* |
| VIL1 | 61 | 96 | 0.217 | 0.013* | 0.06-0.72 | 0.613 | 0.003** |
| ZNRF2 | 78 | 79 | 2.711 | 0.035* | 1.07-6.84 | 0.634 | 0.024* |

**Table 6:** Survival analysis to stratify TNBC patients low-risk and high-risk groups on GSE58812 cohort

| Genes | #G1 | #G2 | HR | p-Value | 95% CI | Concordance | Log(p-value) |
| --- | --- | --- | --- | --- | --- | --- | --- |
| BAIAP3 | 42 | 65 | 0.61 | 0.18 | 0.29-1.26 | 0.55 | 0.19 |
| COL24A1 | 43 | 64 | 0.92 | 0.82 | 0.44-1.92 | 0.51 | 0.82 |
| DHRS7 | 45 | 62 | 0.81 | 0.58 | 0.39-1.69 | 0.54 | 0.58 |
| ETFDH | 58 | 49 | 1.12 | 0.76 | 0.54-2.32 | 0.51 | 0.76 |
| **FRZB** | **46** | **61** | **2.74** | **0.02** | **1.17-6.42** | **0.61** | **0.01** |
| HBG1 | 40 | 67 | 1.05 | 0.90 | 0.49-2.26 | 0.51 | 0.90 |
| IBSP | 56 | 51 | 0.66 | 0.28 | 0.31-1.41 | 0.54 | 0.28 |
| POU4F1 | 40 | 67 | 0.82 | 0.60 | 0.39-1.72 | 0.52 | 0.60 |
| PSAT1 | 54 | 53 | 1.46 | 0.31 | 0.70-3.06 | 0.54 | 0.31 |
| RIOK2 | 61 | 46 | 1.55 | 0.24 | 0.75-3.22 | 0.55 | 0.24 |
| TCEAL4 | 57 | 50 | 0.92 | 0.83 | 0.44-1.92 | 0.52 | 0.83 |
| TCF21 | 46 | 61 | 0.50 | 0.07 | 0.24-1.04 | 0.59 | 0.06 |
| TRPA1 | 38 | 69 | 1.57 | 0.28 | 0.69-3.55 | 0.56 | 0.26 |
| VIL1 | 43 | 64 | 1.40 | 0.39 | 0.65-3.02 | 0.55 | 0.38 |
| **ZNRF2** | **52** | **55** | **2.09** | **0.05** | **0.97-4.50** | **0.59** | **0.05** |

Furthermore, KM survival plots depicted the discrimination of risk groups of LN-TNBC patients (shown in Figure 5).

### Survival Analysis for Clinical Features

Patients’ clinical characteristics, including age, tumor stage, and lymph node status, are considered a primary prognostic indicator for survival in several cancers. Univariate survival analysis was carried out on the clinical features of patients such as age, tumor stage, and lymph node. It was observed that tumor stage and lymph node are important clinical factors with prognostic potential that can stratify LN-TNBC patients into high- and low-risk groups. For example, the tumor stage can significantly stratify risk groups with HR = 11.15 and p-value <0.001, as depicted in Figure 6. Also, lymph nodes significantly stratified risk groups with HR = 4.10 and p-value = 0.002 in TCGA-TNBC patients (Table 7).

**Figure 6:**
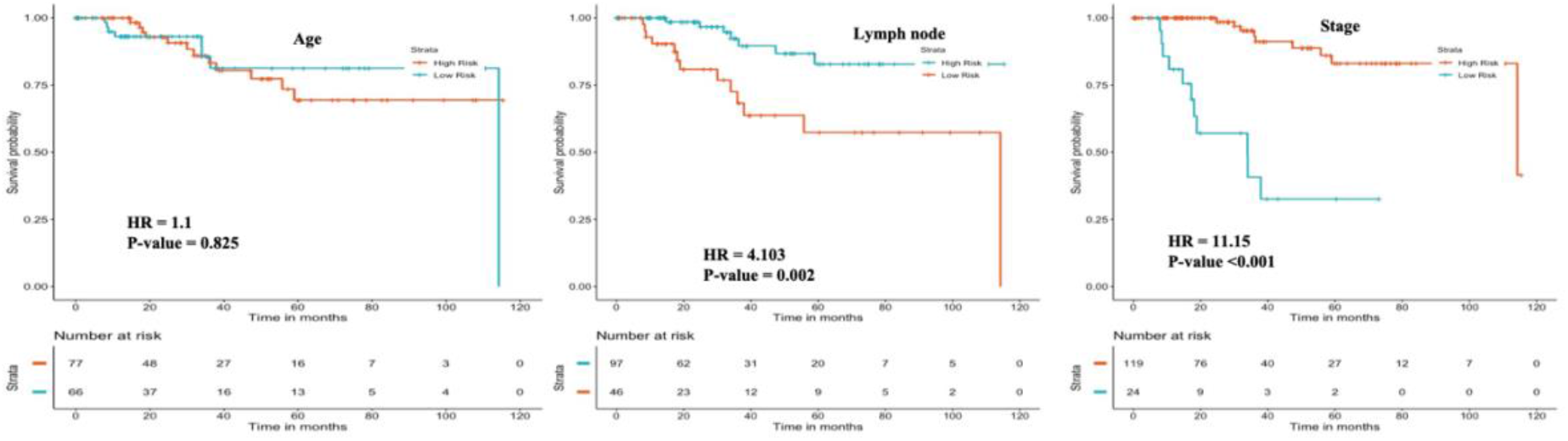
Kaplan Meier survival curves of clinical features (A) Age, (B) LM and (C) Stage

**Table 7:**
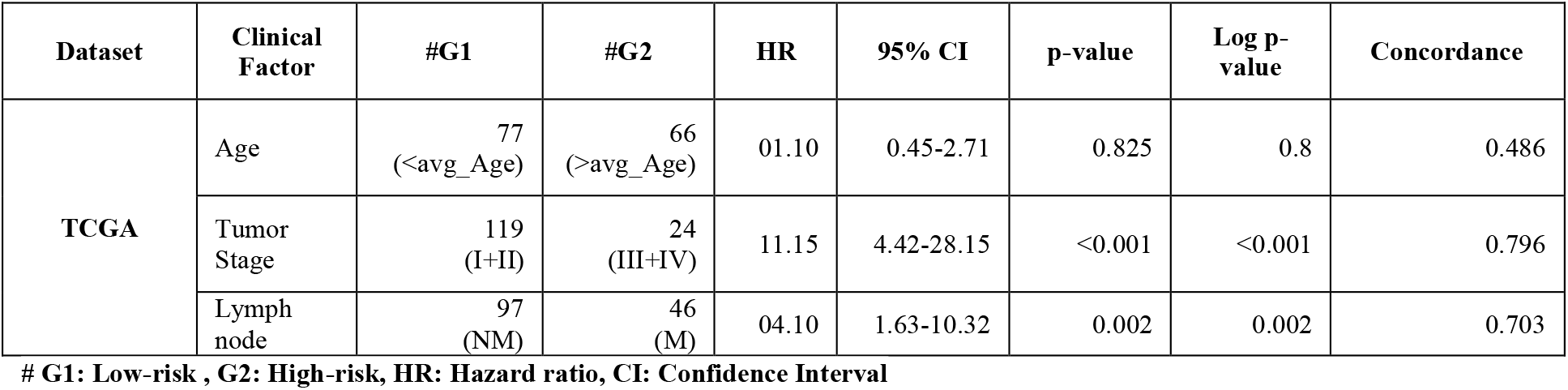
Univariate survival analysis for clinical features of TCGA patients

### Implementation of Webserver

We created a webserver “M_TNBC_Pred” to aid the scientific community working in the field of triple-negative breast cancer research (Prediction server for lymph node metastatic TNBC). In M_TNBC_Pred, two major modules were accomplished: Prediction module and Analysis module based on 5 and 15 genes LNM-TNBC biomarkers. The prediction module allows the users to predict whether the sample is in lymph-node metastatic and non-metastatic using their expression profiles (mRNA expression data) of specific signature genes or biomarkers of TNBC. Here, the user needs to submit log2CPM for a subset of genes or biomarkers. The output result exhibits predicted status like “LN-metastatic or non-metastatic” for each sample. Furthermore, the Analysis module will allow the user to predict whether patient has LN metastatic or non-metastatic based on a single gene. The webserver can be freely accessible at http://webs.iiitd.edu.in/raghava/mtnbcpred/.

## Discussion and Conclusion

Despite the substantial advances in early cancer detection, personalized medicine, and targeted therapy, breast cancer is the second leading cause of demise among females (Siegel et al., 2021b). TNBC has high metastatic tendency and recurrence than other breast cancer subtypes. Where lymph nodes spread is the initial sign of tumor progression and increases the risk of recurrence and mortality (Gonçalves et al., 2018). Tumor-node-metastasis (TNM) staging system for breast cancer, created by the American Joint Committee on Cancer/ International Union Against Cancer describes N classification depend on the number of metastasis lymph nodes. Where N0, lymph node absent, N1, 1–3 metastatic lymph node(s), N2, 4–9 metastatic lymph nodes, and N3, 10 or more metastatic lymph nodes (Byrd, David R., 2010). Thus, it is very important to predict/ diagnose the lymph node metastasis status for TNBC patients. This study analyzed the mRNA-seq expression data of metastatic and non-metastatic LN-TNBC from TCGA to identify gene signatures that act as diagnostic biomarkers.

Firstly, we identified 1010 DEGs (643 upregulated and 367 downregulated genes) with a significant p-value <0.05. Through the logistic-regression method we developed a single gene-based prediction model and found that DHRS7 and ZNRF2 shown the maximum AUC > 0.68. Further, we classify metastatic from non-metastatic LN-TNBC using multiple gene biomarker panels, which incorporate nine upregulated (DHRS7, BAIAP3, ZNRF2, ETFDH, HBG1, RIOK2, TCEAL4, TCF21, FRZB) and six downregulated (POU4F1, COL24A1, TRPA1, IBSP, VIL1, PSAT1). We achieve maximum AUC (0.82 and 0.81) and accuracy (76%, 78.12%) on both training and validation datasets using the GNB classifier. The developed GNB model on 15 genes was used to assess their performance on different independent datasets (Table 3). Our results indicate that these genes may act as a diagnostic marker for the TCGA-TNBC dataset; however, it is quite challenging for other platforms such as Agilent, Illumina, and Affymetrix. Interestingly, these genes show prognostic potential in TNBC patients. For example, FRZB, ZNRF2, and TCEAL4, associated with poor survival, and TRPA1, VIL1, and PSAT1, improves the survival of TNBC patients.

Literature evidence has shown the importance of the few genes found in our analysis and significantly associated with breast cancer metastasis and other cancer types. For instance, FRZB and ZNRF2 were involved in cell growth, increased apoptotic cell death, and cancer metastasis (Araki & Milbrandt, 2003; Shen et al., 2015). The upregulation of FRZB is significantly associated with hepatic metastasis in colon cancer (Shen et al., 2015). ZNRF2 was also widely expressed in nonneural tissues, including the mammary gland, testis, colon, and kidney (Araki & Milbrandt, 2003). Loss of TCEAL4 was observed in anaplastic thyroid cancer (Akaishi et al., 2006). Moreover, the DHRS7 gene regulates the cell cycle in breast cancer cells, can inhibit cell proliferation and expression (Wei-hong, Zhang, 2012). Also, the ectopic expression of BAIAP3 enhances tumorigenic properties in breast and prostate cancer (Palmer et al., 2002).

In conclusion, we have identified the diagnostic potential of the 15 genes in metastasis and non-metastasis TNBC patients with AUC>0.8 on the validation dataset. From survival analysis, we identify three genes ZNRF2 (HR = 2.711, p-value = 0.03), FRZB (HR = 2.395, p-value = 0.05), and TCEAL4 (HR=2.254, p-value = 0.05) are associated with the poor prognosis of metastatic TNBC patients. In addition to this, clinical factors like tumor stage (III/IV) with HR = 11.15, p-value<0.001, and metastatic LN status with HR = 04.10, p-value<0.05 stratify high-risk and low-risk TNBC patients. Thus, clinical factors independently act as major prognostic factors for TNBC patients. Ultimately, we built a freely available webserver, namely M_TNBC_Pred, for predicting metastatic and non-metastatic TNBC using machine learning techniques based on mRNA gene expression data of TNBC patients.

## Authors’ contributions

NLD collected, compiled, and curated the data sets. NLD, AD, SP and GPSR implemented the algorithms and developed the prediction models. NLD, AD, SP, and GPSR analysed the results. SP created the back-end of the web server and AD created the front-end user interface. NLD, AD, SP, and GPSR penned the manuscript. GPSR conceived and coordinated the project, and gave overall supervision to the project. All authors have read and approved the final manuscript.

## Acknowledgements

NLD is thankful to Department of Biotechnology (DBT) for providing Research-Associate fellowship. AD is thankful to the Department of Science and Technology (DST-INSPIRE) and SP is thankful to Department of Biotechnology (DBT) for providing Senior Research fellowships. The authors are thankful to Department of Computational Biology, IIITD New Delhi for infrastructure and facilities.

